# FIB-SEM analysis on three-dimensional structures of growing organelles in wild *Chlorella pyrenoidosa* cells

**DOI:** 10.1101/2022.05.08.491059

**Authors:** Lingchong Feng, Wangbiao Guo, Jiansheng Guo, Xing Zhang, Xiangbo Zou, Mumin Rao, Ji Ye, Cao Kuang, Gongda Chen, Chuangting Chen, Shiwei Qin, Weijuan Yang, Jun Cheng

## Abstract

To clarify dynamic changes of organelle microstructures in *Chlorella pyrenoidosa* cells during photosynthetic growth with CO_2_ fixation, three-dimensional organelle microstructures in three growth periods of meristem, elongation and maturity were quantitatively determined and comprehensively reconstructed with focused ion beam scanning electron microscopy (FIB-SEM). The single round-pancake mitochondria in each cell split into a dumbbell and then into a circular ring, while barycenter distance of mitochondria to chloroplast and nucleus was reduced to 45.5% and 88.3% to strengthen energy transfer, respectively. The single pyrenoid consisting of a large part and another small part in each chloroplast gradually developed to a mature state in which the two parts were nearly equal in size. The nucleolus progressively became larger with euchromatin replication. The number of starch grains gradually increased, but average grain volume remained nearly unchanged.

## 1. Introduction

Photoautotrophic organisms such as higher plants and algae, could convert inorganic carbon from nature into organic carbon via photosynthesis and release oxygen. However, microalgae were 10- to 50-fold more efficient at capturing solar energy for the fixation of carbon dioxide than terrestrial plants (Bhola et al., 2014). The microalgae *Chlorella pyrenoidosa* was a spherical unicellular organism with the beneficial characteristics of a fast growth rate, high photosynthesis rate (Xu et al., 2018; Yu et al., 2019) and the production of many high value-added products (Fernandes et al., 2020; McGee et al., 2020; Patel et al., 2019; Rahman. 2020). The ultrastructure of *C. pyrenoidosa* had been revealed on scanning electron microscopy (SEM) and transmission electron microscopy (TEM) (Andersen et al., 2015; Embleton et al., 2003; Rodenacker et al., 2006; Schulze et al., 2013; Sosik et al., 2007), although only on a two-dimensional XY plane, with more detailed and precise data not able to be acquired in the Z-axis. The 3D structure of microalgal cells had been obtained on laser confocal 3D imaging (Colin et al., 2017), although the low resolution of light microscopy was not enough to reveal organelle microstructural features at the subcellular level.

Focused ion beam scanning electron microscopy (FIB-SEM) was a three-dimensional (3D) imaging technology which can be applied to complete cells or serial slices (Schiel et al., 2011; Uwizeye et al., 2021). FIB-SEM provided the advantages of displaying the shape, number and structure of cellular organelles in 3D, at a high (nanoscale) resolution (Schaffer et al., 2017). With high-pressure freezing (HPF), freeze substitution (FS) and cryogenic resin embedding methods to prepare microalgal samples, it was possible to clarify cell structures in a real state (Wayama et al., 2013). This sample preparation method effectively preserved the original biological information, minimized the damage to cells and maintained the entirety of the cell structure.

Uwizeye et al. (Uwizeye et al., 2021) reported a series of structural images of phytoplankton cells (*Micromonas commoda, Pelagomonas calceolate, Emiliania huxleyi, Galdieria sulphuraria, Phaeodactylum tricornutum* and *Symbiodinium pilosum*), reporting their cell volume and organelle composition from an evolutionary perspective. Wayama et al. (Wayama et al., 2013) reported the structure of *Haematococcus pluvialis* and chlorella cells on three-dimensional transmission electron microscopy (3D-TEM), describing the accumulation of astaxanthin and lipids during the growth of *H. pluvialis*. However, the raw images acquired had a low resolution and their strategy for data segmentation processing was not sufficient, resulting in the cell feature information being incomplete. Flori et al. (Flori et al., 2016) used FIB-SEM to analyze *Phaeodactylum tricornutum* cells, showing that a minimal symbiotic cytoplasm called the periplasmic compartment, exists in the region between the inner and outer membranes of the chloroplast, showing that this region contained the residual cytoplasm of vesicles. However, structural changes in microalgal organelles during their dynamic growth, have not been studied in detail.

In the present study, the advanced imaging technique FIB-SEM was used to visualize the sub-cellular structure of *C. pyrenoidosa*. Then, the volume, surface area and spatial relationships of organelles were quantitatively studied in *C. pyrenoidosa* during different growth periods. The factors responsible for efficient carbon fixation by *C. pyrenoidosa* were established at the organelle level. Furthermore, it was found that the volume ratio of mitochondria to cells remained constant in different periods. It was also observed that the pyrenoid structure was composed of two duplexed parts, the volumes of which remained generally constant during cell growth, with the content of starch grains exhibiting homogeneity.

## 2. Materials and Methods

### 2.1. Cell culture

The microalgae used in this study was *Chlorella pyrenoidosa* (FACHB-9), acquired from the Freshwater Algae Culture Collection at the Institute of Hydrobiology, Chinese Academy of Sciences (China). The *C. pyrenoidosa* strain was maintained in BG11 medium (R. Rippka et al., 1979) in a 300 mL culture bottle, with constant light (~23000 lux) and aeration with 15% CO2 gas (0.1 vvm) at 27 °C for 3 days. The BG11 medium was combined with 20 mmol/L of hepes buffer to maintain stable pH values at 7.2-7.4. The medium was then autoclaved at 121°C for 30 minutes and cooled to room temperature prior to use.

### 2.2 Sample preparation

The high-pressure freezing (HPF) and freeze substitution (FS) reactions were conducted according to a previously reported protocol (Bobik et al., 2014). *C. pyrenoidosa* cultures were centrifuged at 3000 rpm for 5 minutes (ALLEGRAX-15R). Following this, the *C. pyrenoidosa* samples were submerged in a cryoprotectant (1-hexadecene) and carefully loaded into HPF sample carriers (0.1 mm depth, 16770141 Type A, Lecia). The lid (16770142 Type B, Lecia) was submerged in 1-hexadecene and placed on top of the sample carrier. All samples were frozen within several milliseconds under 2100 bar of atmospheric pressure using an EM ICE (Lecia) HPF device (Fig. 1 (B)), then transferred to an EM AFS2 (Lecia) for FS. The sample chamber was filled with ethyl alcohol to maintain a stable temperature. The specimen carrier was transferred to a cap containing 1% uranyl acetate and 2% OsO_4_ in liquid nitrogen, with each cap holding two carriers. The temperature range of FS was minus 90 ° C to 4 ° C (Fig.1 (C)). After FS, samples were washed twice with acetone for 30 min each wash at 4°C, then twice more for 15 min each wash at room temperature. The acetone was then substituted with a 7:3 mixture of acetone: epoxy resin and samples were incubated for 8 hrs. The acetone: epoxy resin solution was replaced by a 3:7 mixture of acetone: epoxy resin and samples were incubated for 12 hrs. Following this, the solution was changed to pure epoxy resin and samples were incubated for 24 hrs, followed by exchange with pure epoxy resin for another 24 hrs. Finally, the polymerized epoxy resin was heated at 60° C in an oven for 48 hrs.

**Figure 1.**
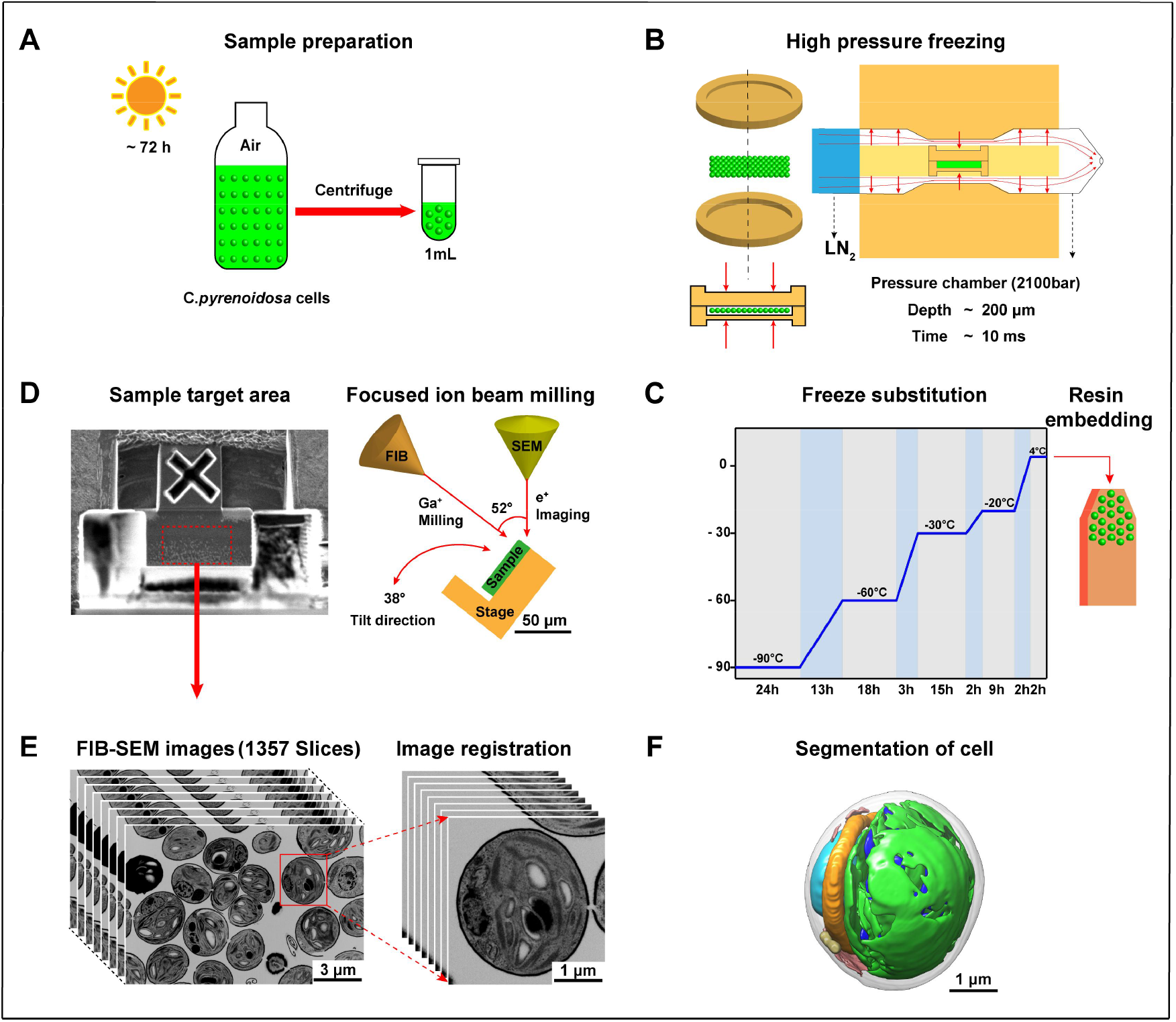
Subcellular visualization process of *C. pyrenoidosa* cells with focused ion beam scanning electron microscopy (FIB-SEM). (A) Microalgal strain culture: *C. pyrenoidosa* cells were incubated in air at light intensity of 23,200 lux for 72 hrs. Samples were centrifuged (3000 rpm for 5 minutes) to obtain 1 mL of microalgal cells slurry. (B) High-pressure freezing of *C. pyrenoidosa:* 10 μL of *C. pyrenoidosa* cells was sampled from 1 mL of microalgal slurry and then transferred to a carrier (0.1 mm depth, Leica). The *C. pyrenoidosa* cells with the carrier were then rapidly frozen in a high-pressure freezer at 2100 bar under liquid nitrogen, with freezing depth of 200 μm and freezing time of approximately 10 ms. (C) Freezing substitution: The frozen samples were treated with osmium acid at −90 to 4°C to complete substitution. The substituted samples were then embedded in resin blocks and fixed in an oven at 60°C for 48 hrs to complete sample preparation process. (D) FIB-SEM process and principles: The FIB was milled by emitting Ga+ and SEM was used for image acquisition by emitting an electron beam, with an angle between FIB and SEM of 53° and samples rotated by 38° after each cycle of completed milling. (E) Image stack of *C. pyrenoidosa* cells (1357 images): Images exhibit a z-axis width of 8 nm and a horizontal field with view width of 17.8 μm. The image segmentation and graphic rendering of intact cells were extracted from raw data by cropping using Amira 6.8 and USCF Chimera software, respectively. (F) Schematic of 3D sub-microstructure of *C. pyrenoidosa* cells.

### 2.3 Data collection by FIB-SEM

After HPF and FS, the samples were embedded in resin blocks and trimmed using a Leica EM trimmer until the black tissue in the block was found. After finding the region of interest, the resin blocks were imaged using a dual beam scanning electron microscope (FIB Helios G3 UC, Thermo Fisher,) (Fig. 1 (D)). Data collection was performed in the serial-surface view mode with a slice thickness of 8 nm and a current of 0.79 nA at 30 keV. Each serial face was imaged using an ICD detector in backscatter mode, with a 2.5 kV acceleration voltage and a current of 0.2 nA. The image store resolution was set to 6144 × 4096 pixels with a dwell time of 2 μs and 2.44 nm per pixel. 1357 slices were acquired and one whole sensilla was contained in the image stack (10.86 μm depth). The horizontal field width of images of *C. pyrenoidosa* were 17.8 μm.

### 2.4 Image processing and segmentation

The FIB-SEM stack of *C. pyrenoidosa* contains 1357 images which include over 20 complete cells. The image sequences of each cell were picked manually and then aligned, cropped, segmented and labeled using Amira 6.8 (Thermo Fisher). Then, the segmented and labeled files were exported for surface rendering and smoothing with USCF Chimera (Pettersen et al., 2004).

The cell volume, surface area, diameter, number and relative coordinate position were measured using the label analysis command in Amira 6.8, which calculated the average, minimum and maximum values for the volume, surface area and diameter of cell components and organelles. The average values were used for calculations. As the shape of *C. pyrenoidosa* was nearly spherical, the equivalent diameter was the spherical diameter. The cell equivalent diameter was calculated according to the average cell volume and the relative surface area was the ratio of surface area to volume.

## 3. Results and discussion

### 3.1. FIB-SEM 3D visualization of the structure of C. pyrenoidosa

*C. pyrenoidosa* cells were selected as a model microalga for analysis, with specimens prepared using high-pressure freezing with subsequent freeze substitution and resin embedding (Bobik et al., 2014). These methods can maximize the visibility of native cell structures, while avoiding sample damage and the formation of structures or debris as a result of chemical fixation. The sample blocks were then subjected to FIB milling (Fig. 1 (D)), followed by SEM generating 2D image data with a Z-axis height of 8 nm. The high resolution allowed subcellular structures to be directly distinguished and classified. Most cells in the field of view had diameters in the range of 2 μm to 4 μm. In total, 1357 2D images were obtained using a sample block with a depth of 10.86 μm (Fig. 1 (E)). To achieve 3D segmentation of *C. pyrenoidosa* cells, the intact cells were selected from the raw image data and Amira 6.8 (Thermo Fisher) was used for image alignment, filtering, noise reduction and contrast enhancement operations. The resulting segmented models were surface rendered using USCF Chimera (Fig. 1 (F)).

The volume segmentation was calculated for 15 intact cells. To discuss the effects of dynamic structural changes in organelles on *C. pyrenoidosa* cells and to explore the interactions between organelles, growth was divided into three periods based on the cell diameter, referred to as the meristem, elongation and mature periods. Přibyl et al. (Přibyl et al., 2012) reported SEM images of chlorella after cultivation for seven days, showing that the chlorella cell diameter was around 2.5 μm in the meristem period, increasing after two days of growth to about 3.0 μm and to 3.5 μm after three days of growth. Based on the measured diameter of *C. pyrenoidosa* cells, cell diameters of 2.5 μm, 3.0 μm and 3.5 μm were defined as the meristem, elongation and mature periods, respectively. Three cells representing each period were selected for 3D reconstruction and their segmented structures were shown in Fig. 2. Results showed that *C. pyrenoidosa* was ellipsoidal in the meristem period, gradually forming a spherical shape in the mature period during cell growth. The cell structure components mainly included the cell wall, nucleus, chloroplast, mitochondria, lipid droplets, vacuole and endoplasmic reticulum. The chloroplast included structures such as the pyrenoid, starch grains, thylakoid membranes and chloroplast stroma. The nucleus mainly consisted of a nucleolus, euchromatin, heterochromatin and the nucleus matrix.

**Figure 2.**
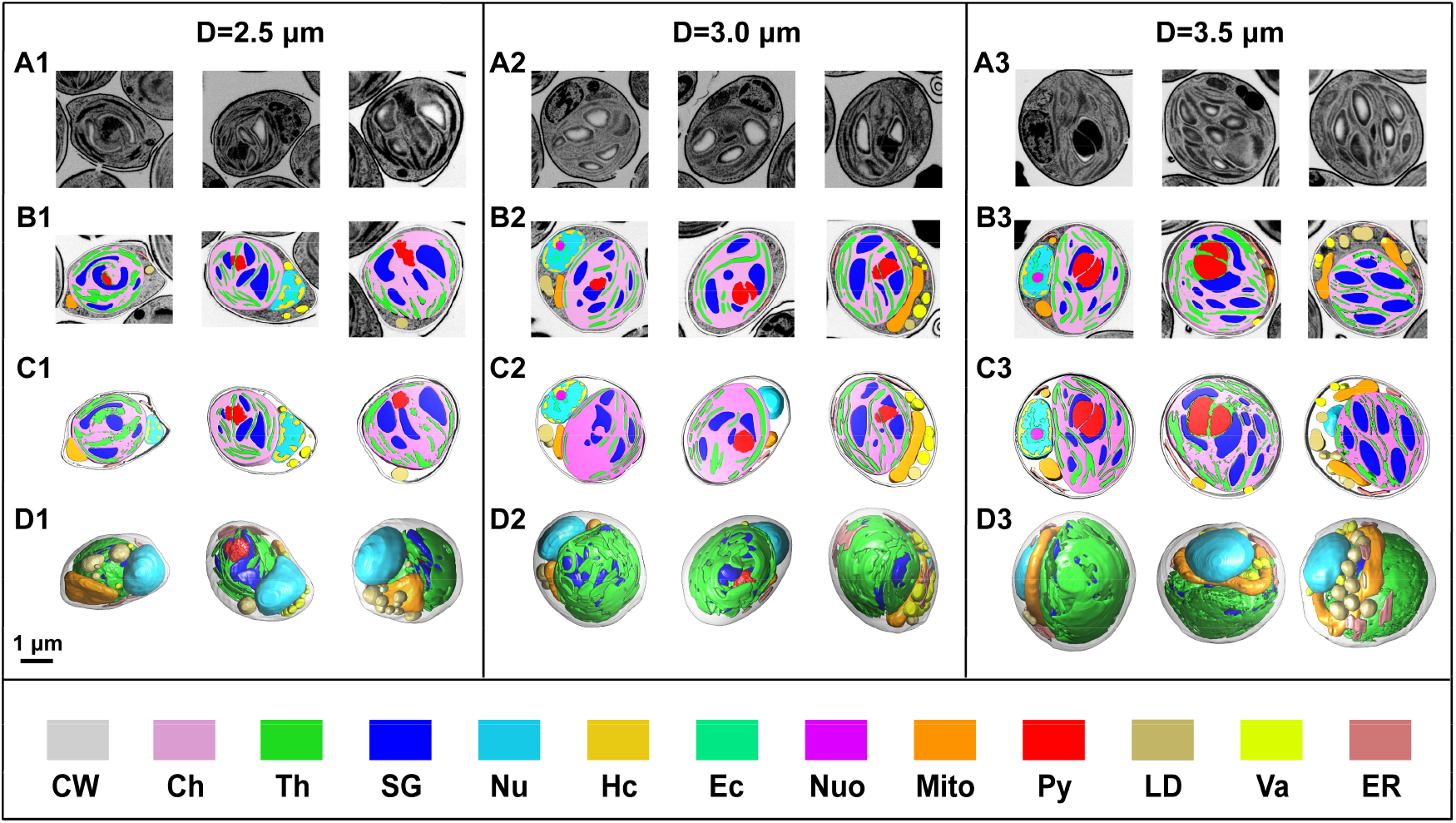
Three-dimensional structure and equivalent spherical diameter (ESD) of *C. pyrenoidosa* cells in three growth periods: meristem period (Diameter=2.5 μm), elongation period (Diameter=3.0 μm) and maturity period (Diameter=3.5 μm). A1, A2 and A3: Original FIB-SEM cross-sections of microalgal cells in different XY planes; B1, B2 and B3: Center-plane segmentation sections of the corresponding cells; C1, C2 and C3: Segmented regions of the corresponding cells at a depth of 160 nm. D1, D2 and D3: Three-dimensional segmented structures of the corresponding cells. Scale bar: 1 μm. **Note:** cell wall (CW); chloroplast (Ch); thylakoid (Th); starch grain (SG); pyrenoid (Py); vacuole (Va); endoplasmic reticulum (ER); mitochondrion (Mito); nucleus (Nu); lipid droplet (LD); nucleolus (Nuo); euchromatin (Ec); heterochromatin (Hc).

By comparing the 3D structure of *C. pyrenoidosa* organelles, it was found that only one mitochondria was present in the whole cell, unlike higher plant cells which contained multiple mitochondria. The mitochondria in *C. pyrenoidosa* was in the middle of the chloroplast and the nucleus, which was convenient for the supply of ATP to each organelle. In addition, the shape of mitochondria was dynamic, changing with cell growth from a round-pancake structure in the meristem period to a dumbbell structure in the elongation period, finally forming a circular ring around the interior of the cell in the mature period. The pyrenoid in the chloroplast was found to be a duplex structure surrounded by starch granules, containing a high abundance of rubisco enzymes (ribulose-1,5-bisphosphate carboxylase/oxygenase) for the capture of carbon dioxide. Starch grains were formed in gaps of the lamellar structure of the thylakoid membrane, then distributed in the chloroplast stroma. The nucleus was located at the edge of the cell, near the cell membrane. The heterochromatin inside the nucleus was evenly distributed on the inner membrane of the nucleus and the euchromatin was free in the nucleolar matrix. The nucleolus was a uniform spherical structure including a high abundance of rRNA, rDNA and ribosomal proteins. The endoplasmic reticulum, vacuole and lipid droplets were irregularly distributed around the mitochondria and nucleus. Because of the resolution limit, the vesicle structure of the Golgi apparatus could not be accurately acquired from images and thus, its 3D structure could not be established. In the raw data acquired by FIB-SEM, it was observed that the cell wall was broken (Fig. 2 (A2)), which might be due to 1-hexadecene (used as a cryoprotectant during the high-pressure freezing process) not being uniformly mixed with cells. Subsequently, the mechanical strength of the cell wall might be unable to withstand the rapid increase in pressure (2100 bar), causing rupturing of the cell wall surface. Cellulose and glycoproteins had been shown to be the main components of the chlorella cell wall (Domozych et al., 2012), which were used to support and maintain the shape of cells, ensuring that damage to the cell wall does not affect internal structural features.

### 3.2. Analysis of the occupancy of organelles in C. pyrenoidosa cells in different cell growth periods

The 3D segmentation models of *C. pyrenoidosa* in different growth periods were presented in Fig. 3 (A), showing that there were obvious differences in the number, location, volume and surface area of organelles in each growth period. The volume ratios of organelles to cells in the meristem, elongation and mature periods were shown in Fig. 3 (B). The results showed that *C. pyrenoidosa* contained only one large chloroplast, which occupied more than half of the cell volume, while the chloroplast of higher plants had been reported to occupy only 40% of the cytoplasm. Compared to higher plant cells, the chloroplast in *C. pyrenoidosa* contained a larger thylakoid membrane area, providing more photosynthetic pigments for the capture of light energy, greatly improving the efficiency of photosynthesis. As the cell grew, the percentage volume of the chloroplast and its major internal structures became gradually larger, with the increased volume ratio of the thylakoid and pyrenoid to the whole cell, resulting in a higher photosynthetic efficiency. The photosynthetic capability of *C. pyrenoidosa* was enhanced from the meristem period to the mature period, with the accumulation of organic matter such as starch grains also increased. Lipid droplets exhibited the largest volume ratio during the meristem period, with the pre-divided cells containing more lipid droplets and the newly formed cells containing lipid droplets supplied from the mother cells. At this point, the chloroplast was not in a mature state and the organic matter accumulated by photosynthesis was not sufficient to meet their requirements for growth. Therefore, to support the initial rapid growth of cells, lipid droplets were metabolized, and the proportion decreased. In contrast, when cells grew to maturity, their photosynthetic capacity increased and the endoplasmic reticulum was developed, and lipid droplets were synthesized, resulting in a gradual increase in their proportion.

**Figure 3.**
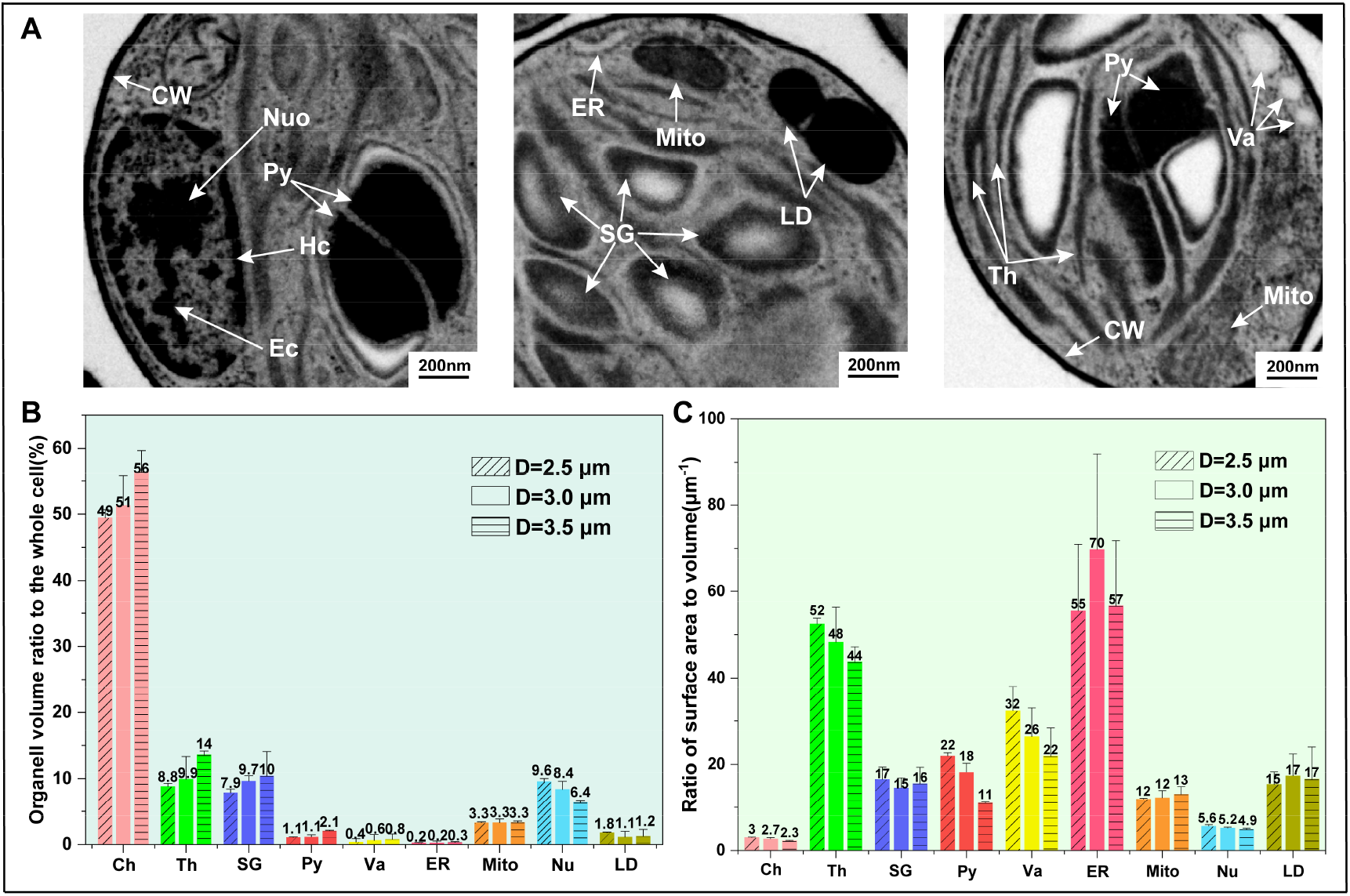
Classification of organelles in XY plane of *C. pyrenoidosa* cells according to their volume ratios to overall cell and relative surface areas (n= ≥3). (A) Major cell structures: cell wall (CW), chloroplast (Ch), thylakoid (Th), starch grain (SG), pyrenoid (Py), vacuole (Va), endoplasmic reticulum (ER), mitochondria (Mito), nucleus (Nu), lipid droplet (LD), nucleolus (Nuo), euchromatin (Ec), heterochromatin (Hc); (B) Volume ratio of each organelle to overall cell volume in three cell growth periods: meristem period (Diameter=2.5 μm), elongation period (Diameter=3.0 μm) and maturity period (Diameter=3.5 μm); (C) Relative surface area: Ratio of surface area to volume for each organelle. Scale bar: 200 nm.

Fig. 3 (B) showed the relative surface area (μm^-1^), which indicated the ratio of organelle surface area to the volume of *C. pyrenoidosa* cells during all three growth periods. These results indicated that the relative material transportation efficiency inside the chloroplast decreased gradually with cell growth. In the meristem period, the relatively smaller volume of the thylakoid membrane was dispersed into lamellar structures, forming a larger light trapping area and thus, increasing the light absorption capacity of the cell. The surface area to volume ratio of *C. pyrenoidosa* cells decreased due to cells had a large relative surface area in the meristem period, resulting in microalgae being more efficient at transportation and cell metabolism, allowing rapid information transfer. During microalgal growth, cells needed to obtain more nutrients from the external environment, although the increase in cell surface area was relatively less, leading to cell growth becoming gradually limited.

### 3.3. The distance between the barycenter of mitochondria, the chloroplast and nucleus became smaller, improving the efficiency of energy supply for cell growth

The 3D structure of cells reconstructed from FIB-SEM image stacking, was used to establish the overall topology of cells and provide in-depth information about the interactions between organelles during cell growth. The structural morphology of mitochondria reflected its functionality, maintaining normal cell physiological activities and metabolism. During the three periods of cell growth, significant differences were observed in the mitochondrial topology (Fig. 4 (A1, A2, A3)), exhibiting a round-pancake shape in the meristem period, splitting into a dumbbell structure in the elongation period and fusing to form a circular ring in the mature period. This process of morphological transformation during mitochondrial fission and fusion, was crucial for effective environmental adaptation and the rapid growth of phytoplankton such as microalgae (Tilokani et al., 2018). Furthermore, it was also found that the relative positions of the mitochondria, chloroplast and nucleus remained unchanged, with the chloroplast and nucleus located on both sides of the mitochondria, as shown in Fig. 4 (B1, B2, B3). However, the distance between the barycenter of mitochondria to the chloroplast and nucleus barycenter was reduced (Fig. 4 (C)), described as distances d1 and d2, respectively (Fig. 4 (B)). When *C. pyrenoidosa* cells were in the meristem period, d1 and d2 exhibited maximum values of 1.34 μm and 1.11 μm, respectively, reducing after cells reached maturity to 0.61 μm and 0.98 μm, indicating a shortening by 45.5% and 88.3%, respectively. The reduced distance between organelle barycenters resulted in less energy required for material exchange. In particular, the large reduction in d1 value was beneficial for mitochondria, allowing adequate energy to be provided for vital activities in the nucleus, such as rRNA transcription and chromatin replication, promoting cell growth overall.

**Figure 4.**
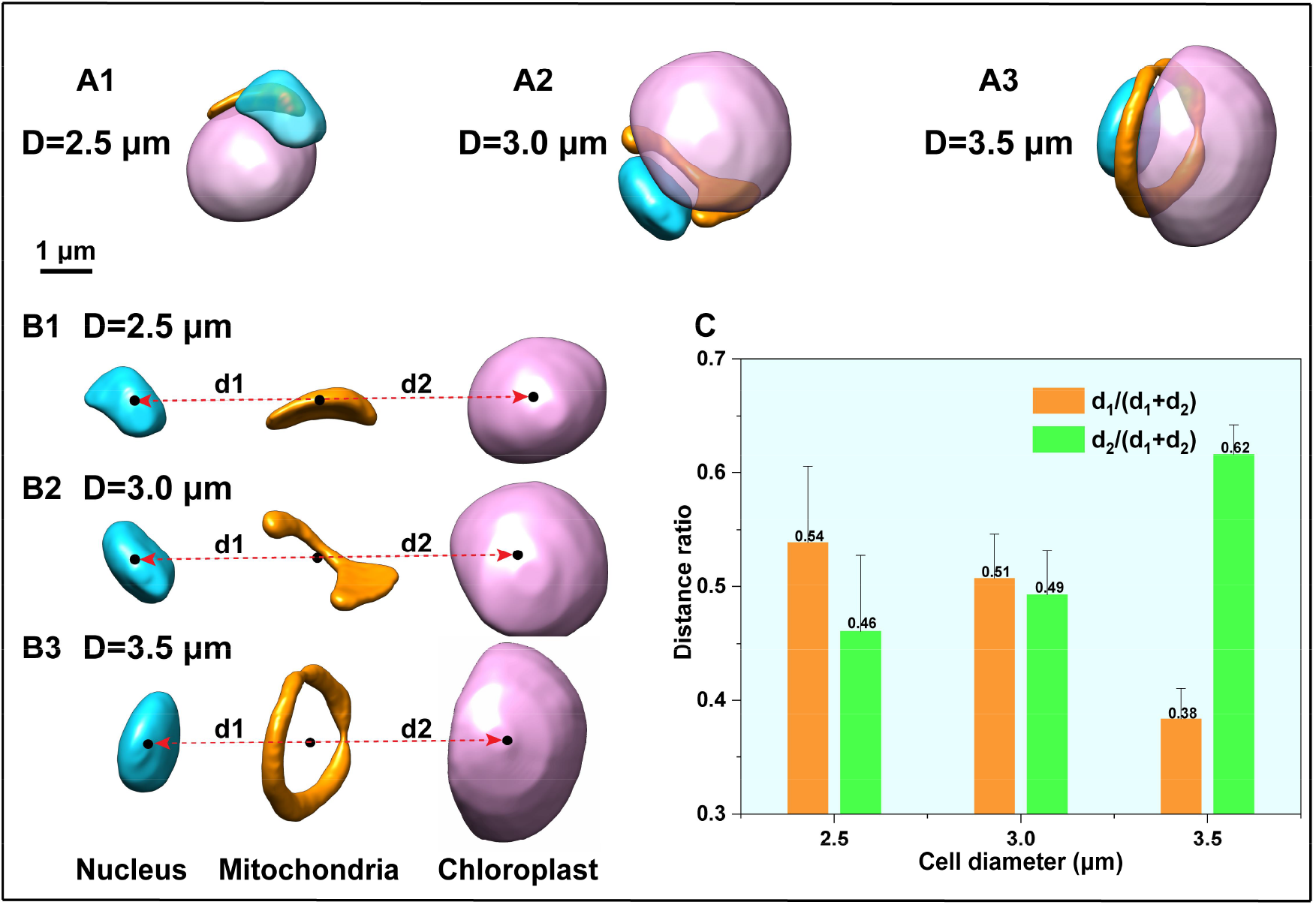
Morphological structure and position analysis on nucleus, mitochondria and chloroplasts of *C. pyrenoidosa* cells in three growth periods: meristem period (Diameter=2.5 μm), elongation period (Diameter=3.0 μm) and maturity period (Diameter=3.5 μm). A1, A2 and A3: Morphological changes of mitochondria from round pancake shape to dumbbell type with center splitting and further to circular ring type. B1, B2 and B3: Changes in relative position of mitochondria to nucleus and chloroplasts. The nucleus and chloroplasts reside on either side of mitochondria, with certain distances (d1 and d2) between mitochondria barycenter and chloroplast barycenter as well as nucleus barycenter. C: Distance changes in nucleus barycenter and chloroplasts barycenter relative to mitochondria barycenter. Scale bar: 1 μm.

In this study, the mitochondria-to-cell volume ratio was found to remain stable at approximately 3.31%, while the chloroplast-to-cell volume ratio increased and the nucleus-to-cell volume ratio reduced as cells grew (Fig. 3 (B)). This indicated that the mitochondrial volume consistently maintained a positive relationship with the cell volume, which has major significance in the study of the coordinated interactions between mitochondria and other cellular structures in phytoplankton. Flori et al. (Flori et al., 2017) and Mueller-Schuessele et al. (Mueller-Schuessele et al., 2018) reported that mitochondria-chloroplast interactions were likely to originate from physical dependence, with common contact points and/or surfaces. Engel et al. (Engel et al., 2015) found that in diatoms, structural interactions between mitochondria and chloroplasts ensured the sharing of reducing equivalents and ATP, facilitating the light assimilation of CO_2_. Mitochondria and chloroplasts exhibited a very close physical relationship, allowing energetic interactions between the two organelles to be possible. In the present study, although mutual contact was not observed directly in the 3D structure, this may be due to the small size of the contact area, with some details not identifiable during post-processing. However, the results of the present study showed that the distance between the mitochondria and chloroplast barycenter (d2) was reduced, which supported the proposed energetic interaction between these organelles. Results also showed that the nucleus was significantly closer to the mitochondria than to the chloroplast, as the nucleus was metabolically advanced when cell growth reached a mature stage. The nucleus generated large amounts of RNA for transcription and the translation of proteins by ribosomes in the cytoplasm, with this process requiring high levels of ATP. Moreover, chloroplast light reactions produced enough ATP for reactions to continue in the dark and therefore, the nucleus was more dependent on mitochondria than the chloroplast.

### 3.4. Enhanced information exchange between the nucleus and chloroplast promoted cell growth and carbon fixation

The nucleus was the regulatory center of cell genetics and metabolism. Therefore, the 3D structure of the nucleus and its internal genetic material was reconstructed and quantitatively analyzed (Fig. 5 (A1-B1, A2-B2, A3-B3)). Results showed that the nucleus was located at the edge of the cell near the cell membrane, while the volume ratio (N/C volume ratio) and relative surface area of the nucleus to the perinucleus cytoplasm gradually decreased with cell growth. The volume ratio of the nucleus to the cytoplasm reflected the strength of cell viability, especially in the meristem period, which exhibited a higher N/C ratio value and was more dynamic (Sokoloff, 1922). Following this, the relative surface area of the nucleus also decreased as the cell volume become larger, indicating a relative reduction in the material transfer efficiency. This suggested that *C. pyrenoidosa* cells in the meristem period were more active metabolically, resulting in the nucleus needing a larger relative surface area, providing more nuclear pores to transmit biomolecules to the cytoplasm, such as RNA and proteins.

**Figure 5.**
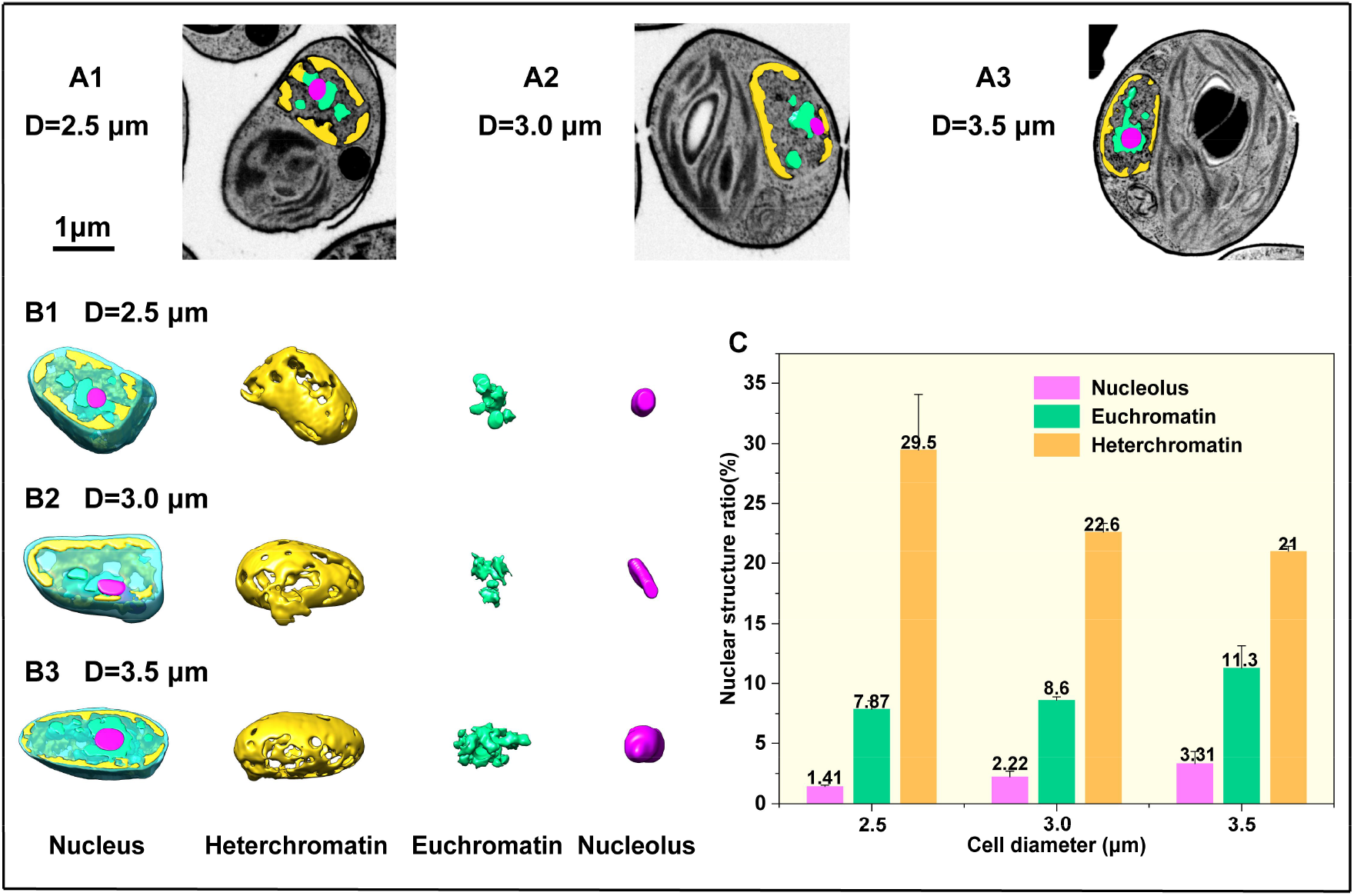
Morphological and structural analysis on *C. pyrenoidosa* cells in three growth periods: meristem period (Diameter=2.5 μm), elongation period (Diameter=3.0 μm) and maturity period (Diameter=3.5 μm). A1, A2 and A3: FIB-SEM cross-sections of nucleus structures; B1, B2 and B3: Three-dimensional structures of nucleus, heterochromatin, euchromatin and nucleolus; C: Proportion variations of nucleolus, euchromatin and heterochromatin in total nucleus area. Scale bar: 1 μm.

Chromatin, the nucleolus and the nucleus matrix were the primary structures of the nucleus, in which chromatin was divided into heterochromatin (Fig. 5 (D1, D2, D3)) and euchromatin (Fig. 5 (E1, E2, E3)). This study focused on the regulatory effect of changes in the volume of chromatin and nucleolus during cell growth, with less focus was paid to the nucleus matrix. The nucleolus and chromatin were quantitatively studied, finding the nucleolus to nucleus volume ratio to be 1.41% in the meristem period, 2.22% in the elongation period and 3.31% in the maturity period. The nucleolus volume ratio enlarged 1.35-fold from the meristem to mature period. The euchromatin volume ratio was 7.87%, 8.6%, and 11.3% in the meristem, elongation and maturity periods of cell growth, respectively, with the relative volume ratio of euchromatin in the mature period being 1.44-fold higher than in the meristem period. However, the volume ratio of heterochromatin exhibited an opposite trend, with a maximum ratio of 29.5% observed in the meristem period, reducing to 22.6% and 21% in the elongation and mature periods, respectively, showing a decrease in relative volume ratio by 28.8% from the meristem to the mature periods. The relative volume ratio of the nucleolus became larger because *C. pyrenoidosa* cells needed to produce more proteins during the rapid growth period. Ribosomes were the first active organelle in the translation and synthesis of proteins and thus, a high abundance of ribosomes were synthesized to meet the demands. Ribosomes were composed of rRNA and ribosomal proteins, with the main function of the nucleolus being the synthesis of rRNA and the formation of ribosomes. With gradual enlargement of the cell volume, the nucleolus volume also enlarged to satisfy the protein requirements for cell growth. It was found that the euchromatin volume ratio also increased as the volume of the cell increased. Euchromatin replicated during the S-phase of cell cycles, at which point gene density was higher and most genes can be expressed, with enough transcriptional activity to translate and produce proteins. Therefore, euchromatin replicated preferentially during the growth of microalgal cells. Although the relative volume of heterochromatin decreased with cell growth, its absolute volume (0.27 μm^3^, 0.27 μm^3^ and 0.29 μm^3^, in the meristem, elongation and mature periods, respectively) did not decline. This may be because heterochromatin was a tightly assembled structure distributed in the edge of nucleus, with a high histone encapsulation rate and a high degree of double helicization, in which most genes did not have genetic activity and thus, cannot transcribe proteins. In the cell cycle, heterochromatin replicated later than euchromatin and therefore, its relative volume decreased as the cell grows, with the results of the present study indicating that heterochromatin gradually started to replicate when cells were in the mature period.

In eukaryotic algal cells, chloroplasts and mitochondria contained genetic DNA material, which can encode the transcriptional synthesis of proteins. For example, the chloroplast of *Chlamydomonas reinhardtii* had been shown to contain 72 protein-coding genes (Maul et al., 2002), while eight protein-coding genes were retained in the mitochondria (GenBank Accession No. CRU03843). Chloroplasts contained about 3000 different proteins (Abdallah et al., 2000) and they were not able to meet the demands of photosynthesis using only their own encoded protein genes, with most proteins encoded by the nucleus, synthesized in the cytoplasmic stroma and then transported to the chloroplast. Accordingly, the volume ratio of genetic material to the nucleolus reflected the physiological state of the cell, with a larger volume ratio also enhancing the photosynthetic carbon sequestration capacity of the chloroplast, promoting further cell growth and development.

### 3.5. Duplex structure of the pyrenoid enhanced dark reaction rates and material synthesis

The pyrenoid structure within the chloroplast exhibited a unique duplex shape (Fig. 6 (C1, C2, C3)), which was composed of two parts, one of which was larger than the other, with the volume of both parts found not to grow simultaneously. A tube similar to the thylakoid membrane was observed between the two layers of the duplex structure, communicating with the pyrenoid and thylakoid (Fig. 6 (B1)). The 3D structures of the small volume pyrenoid (Py_S_) and the large volume pyrenoid (Py_L_) were quantitatively analyzed, revealing that the volume ratio of Py_S_ to pyrenoid increased with cell growth, with volume ratio values in the meristem, elongation and mature periods of 22.1%, 38.5% and 47.2%, respectively. In contrast, the volume ratio of Py_L_ to pyrenoid decreased with growth, with volume ratio values in the meristem, elongation and mature periods of 77.9%, 61.5% and 52.8%, respectively (Fig. 6 (D)). During the early phase of *C. pyrenoidosa* growth, the volume of Py_S_ and Py_L_ were considerably different and with cell growth, the Py_S_ growth rate was significantly faster than that of Py_L_, resulting in the volume of the two parts gradually becoming similar during the mature period.

**Figure 6.**
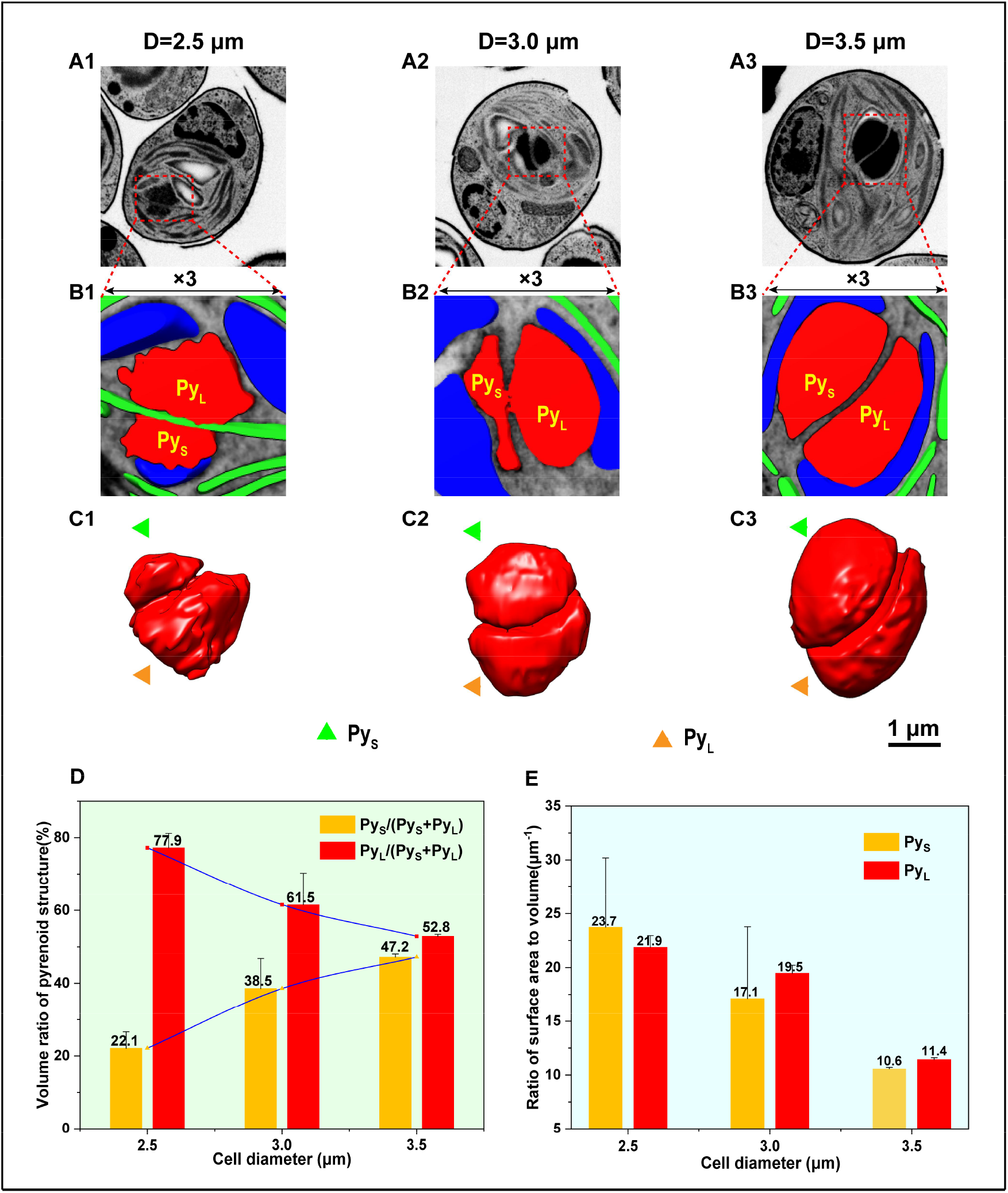
Morphological structure analysis on pyrenoids of *C. pyrenoidosa* cells in three growth periods: meristem period (Diameter=2.5 μm), elongation period (Diameter=3.0 μm) and maturity period (Diameter=3.5 μm). A1, A2 and A3: FIB-SEM mapping of original *C. pyrenoidosa* cells with pyrenoids as target regions (in red dashed boxes). B1, B2 and B3: FIB-SEM 3D analysis on the corresponding segmented sections of target pyrenoid regions at three-fold magnification. C1, C2 and C3: 3D topology of pyrenoid in target regions, consisting of two parts: PyL and PyS in duplex structures. D: Relative volume ratios of PyL and PyS to the whole pyrenoid. E: Relative surface areas of PyL and PyS to pyrenoid volume. Scale bar: 1 μm.

The relative surface area of Py_S_ and Py_L_ both decreased with cell growth and therefore, the pyrenoid relative surface area also decreased, while the Py_S_ relative surface area was higher than that of Py_L_ when cells were in the meristem period (Fig. 6 (E)). This may indicate a supportive relationship or affiliation, in which one side passes Rubisco enzymes, carbonic anhydrase, ATP and the other materials required for reactions to occur under dark conditions via a duct similar to the thylakoid membrane (Engel et al., 2015; Rochaix, 2017). With gradual cell growth, the rate of dark condition reactions in photosynthesis was gradually enhanced, increasing the accumulation of organic matter and resulting in the total volume of the pyrenoid becoming progressively larger. In contrast, Py_S_ initially had a larger volume growth rate than Py_L_ due to the accumulation of organic matter and the supply of materials from Py_L_, with its surface area to volume ratio and relative growth rate gradually decreasing. The size of the pyrenoid was limited by its surrounding starch sheath and it was not able to grow freely, resulting in the volume ratio of Py_S_ to Py_L_ becoming close to 1 in the mature period. Moreover, the pyrenoid was a liquid-like organelle (Itakura et al., 2019; Freeman Rosenzweig et al., 2017; Mackinder et al., 2017), surrounded by starch sheath. Under natural conditions, *C. pyrenoidosa* fixes CO_2_ through the CCM mechanism, which occured on Rubisco, with the major Rubisco enzymes existing in the pyrenoid. In the CCM theory, when the external CO_2_ concentration was low, microalgal cells useed HCO^3-^ as an additional carbon source to enter the cell by active transport and then be fixed by the pyrenoid to achieve high photosynthetic efficiency. The surface area to volume ratio of the pyrenoid was an important parameter for the CO_2_ absorption capacity of *C. pyrenoidosa*. In the early period of cell growth, the pyrenoid was flattened, increasing the contact sites with CO_2_ by forming the largest possible surface area, while also maximizing the contact and transfer of substances with the thylakoid membrane to promote reactions under dark conditions and the rapid growth of cells. In the later phase of cell growth, substance accumulation and enzymes expression increased, leading to gradual enlargement of Py_S_. The rate of cellular material synthesis was reduced relatively, with growth gradually stabilizing, allowing the Py_S_ and Py_L_ structures of the pyrenoid to meet the demands of dark condition reactions.

### 3.6. Starch grains were synthesized and converted maintaining a stable average volume and energy supply

In microalgal cells, starch grains were polymerized from glucose produced by photosynthesis and located in the chloroplast stroma. Starch grains were separated in sequence by the lamellar structure of the thylakoid membrane (Fig. 7 (B1, B2, B3)), with grains exhibiting an ellipsoidal shape bearing two sharp sides (Fig. 7 (C1, C2, C3)). Starch grains closer to the pyrenoid were found to have a larger volume, with the statistical average number and volume of starch grains analyzed in different growth periods. At the beginning of cell growth (the meristem period), the number of starch grains in individual cells was low, with the number of grains increasing with cell growth. It was also found that the maximum volume of individual starch grain had a threshold value, with the threshold value becoming larger with cell growth were 0.25μm^3^, 0.29μm^3^, 0.38μm^3^, respectively (Fig. 7(D)). However, the average volume of a single starch grain within *C. pyrenoidosa* cells remained constant at around 0.12 μm^3^ with ongoing cell growth (Fig. 7(E)).

**Figure 7.**
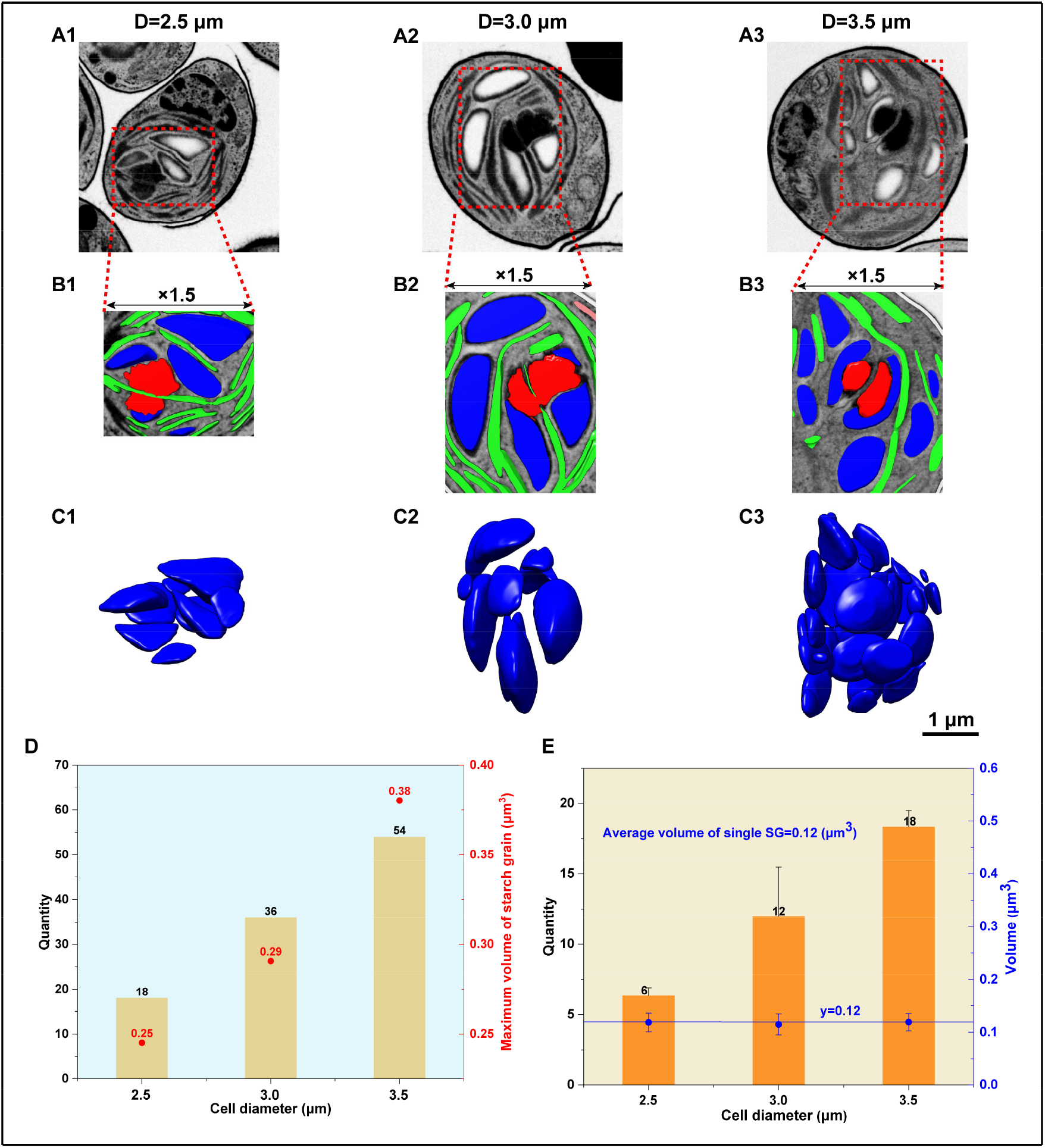
Number and volume analysis on starch grains of *C. pyrenoidosa* cells in three growth periods: meristem period (Diameter=2.5 μm), elongation period (Diameter=3.0 μm) and maturity period (Diameter=3.5 μm). A1, A2 and A3: FIB-SEM mapping of original *C. pyrenoidosa* cells with starch grains as target regions (in red dashed boxes). B1, B2 and B3: FIB-SEM 3D analysis on the corresponding cross sections of target regions at 1.5-fold magnification. C1, C2 and C3: 3D topology of starch grains in target regions. (D) Volume distributions of starch grains. (E) Number of starch grains and average volume of individual starch grains. Scale bar: 1 μm.

As reported previously, the volume ratio of pyrenoids and starch grains to the whole cell gradually became larger with ongoing growth, indicating that pyrenoids play a positive role in the synthesis and accumulation of starch grains, which was consistent with the conclusions of Meyer et al. (Meyer et al., 2017) Uwizeye et al. (Uwizeye et al., 2021)proposed that the first step in starch synthesis (ADP-glucose pyrophosphorylase formation of ADP-glucose) was stimulated by 3-phosphoglyceric acid (3PGA), which was a direct product of CO_2_ fixation by Rubisco enzymes in the Calvin-Benson cycle. As discussed previously, the average volume of starch grains was found to be about 0.12 μm^3^. It may be proposed that the accumulation and consumption of starch grains and their conversion to lipids during photosynthesis was a dynamic process, with the average volume of starch grains influenced by the accumulation of dark condition reaction materials. Starch grains were produced in the chloroplast stroma between the gaps of the thylakoid membrane and if the volume of starch grains becomes too large, the distance required for material exchange between membranes was increased, negatively affecting photosynthesis. Furthermore, starch grains were stored in the starch sheath, limiting the maximum possible volume of each grain. Svihus et al. (Svihus et al., 2005) and Tester et al. (Tester et al., 2006) showed that the size, shape, specific surface area and composition of starch grains were all important factors affecting the hydrolysis of starch. Vasanthan et al. (Vasanthan et al., 1996) extracted different particle size starch grains from barley and found that small grains were easier hydrolyzed by β-amylase. Larger starch grains had a smaller relative surface area which was less conductive to enzyme catabolism. However, as microalgal cells grow, the activity and expression of starch hydrolases such as α-amylase, β-amylase and starch phosphorylase became larger, making microalgal cells more capable of hydrolyzing starch grains and thus, the maximum volume of starch grains was increased. The average volume of starch grains remained generally constant across all three growth periods, indicating that during the meristem and elongation periods, starch grains accumulate for conversion to ATP and proteins synthesis. After microalgal cells were in the mature period, the production of starch grains increased and the consumption rate by cells became faster. In addition to proteins and ATP, starch grains were also converted into lipids that were stored in cells with a higher energy density so that after cell division, cells in the meristem stage consume lipids, providing energy for cell growth when photosynthesis rates were low.

## 4. Conclusions

Quantitative analysis of the three-dimensional structure of organelles in *C. pyrenoidosa* on FIB-SEM, clarified the interactions between organelles in dynamically growing cells, with changes in their positional relationships effectively promoting carbon fixation by photosynthesis. Changes in the volumes of the nucleolus and chromatin indicated variations in cellular activity, regulating gene expression to influence cellular behavior and accomplish vital *C. pyrenoidosa* cell activities such as metabolism, growth and reproduction. The duplex structure of the pyrenoid promoted reactions under dark conditions and the production and accumulation of starch grains. The large chloroplast and mitochondria structures of *C. pyrenoidosa* reflected the higher photosynthetic efficiency of chlorella at the organelle level than terrestrial higher plants. In future studies, it will be necessary to analyze the mechanism of action at the protein molecular level for typical organelles of *C. pyrenoidosa*.

**Table 1:** Volume of 15 chlorella cells(μm^3^)

| Volume of 15 chlorella cells(μm³) |
| --- |
| 9.3 |
| 9.6 |
| 9.9 |
| 12.1 |
| 12.2 |
| 12.4 |
| 14.1 |
| 14.5 |
| 16.0 |
| 17.2 |
| 19.1 |
| 20.5 |
| 20.9 |
| 21.9 |
| 22.3 |

**Table 2:** Volume of cell organelles(μm^3^)

| Volume of cell organelles(μm³) |  |  |  |  |  |  |  |  |  |  |  |
| --- | --- | --- | --- | --- | --- | --- | --- | --- | --- | --- | --- |
| Select of 9 chlorella cells(μm³) | Number | Cell | Ch | Th | SG | Py | Nu | Mito | LD | Va | ER |
| Meristem | 1 | 9.3 | 4.62 | 0.77 | 0.72 | 0.1 | 0.88 | 0.3 | 0.17 | 0.09 | 0.026 |
|  | 2 | 9.55 | 4.83 | 0.88 | 0.81 | 0.11 | 0.96 | 0.31 | 0.16 | 0.006 | 0.015 |
|  | 3 | 9.92 | 4.78 | 0.89 | 0.74 | 0.1 | 0.91 | 0.34 | 0.18 | 0.009 | 0.027 |
| Elongation | 4 | 12.1 | 5.7 | 1.02 | 1.06 | 0.16 | 1.16 | 0.38 | 0.2 | 0.016 | 0.002 |
|  | 5 | 14.1 | 7.14 | 1.95 | 1.4 | 0.11 | 1.19 | 0.55 | 0.02 | 0.23 | 0.08 |
|  | 6 | 17.2 | 9.67 | 1.29 | 1.77 | 0.23 | 1.23 | 0.5 | 0.27 | 0.01 | 0.016 |
| Mature | 7 | 20.9 | 11.3 | 2.72 | 1.35 | 0.45 | 1.4 | 0.72 | 0.017 | 0.36 | 0.04 |
|  | 8 | 21.9 | 12.1 | 2.98 | 2.98 | 0.44 | 1.39 | 0.76 | 0.38 | 0.06 | 0.07 |
|  | 9 | 22.3 | 13.4 | 3.16 | 2.51 | 0.46 | 1.34 | 0.7 | 0.43 | 0.07 | 0.1 |

**Table 3:** Area of cell organelles(μm^2^)

| Area of cell organelles(μm²) |  |  |  |  |  |  |  |  |  |  |  |
| --- | --- | --- | --- | --- | --- | --- | --- | --- | --- | --- | --- |
| Select of 9 chlorella cells(μm²) | Area(μm²) | Cell | Ch | Th | SG | Py | Nu | Mito | LD | Va | ER |
| Meristem | 1 | 23 | 14.4 | 39.4 | 11.3 | 2.21 | 5.28 | 3.63 | 2.39 | 2.33 | 1.89 |
|  | 2 | 23.4 | 14.2 | 46.1 | 11.6 | 2.33 | 5.1 | 3.66 | 2.99 | 0.22 | 0.64 |
|  | 3 | 23.9 | 14.3 | 47.9 | 14.6 | 2.26 | 5.03 | 4.1 | 2.42 | 0.31 | 1.38 |
| Elongation | 4 | 33 | 17.1 | 43.3 | 15.9 | 2.69 | 5.96 | 5.03 | 3.14 | 0.47 | 0.19 |
|  | 5 | 29.5 | 18.7 | 87.8 | 17 | 2.26 | 6.46 | 5.73 | 0.46 | 4.32 | 4.3 |
|  | 6 | 42.6 | 23.8 | 74.1 | 29.5 | 3.98 | 6.15 | 6.63 | 3.63 | 0.31 | 0.97 |
| Mature | 7 | 46.4 | 26.4 | 109.6 | 26.7 | 4.89 | 6.57 | 9.76 | 0.42 | 5.36 | 2.96 |
|  | 8 | 47.8 | 26.6 | 130.6 | 37.5 | 4.84 | 6.94 | 11.1 | 5.64 | 1.65 | 3.24 |
|  | 9 | 48.6 | 29.9 | 149.1 | 36.2 | 5.17 | 6.83 | 7.88 | 4.42 | 1.64 | 4.97 |

**Table 4:** Volume and area of nucleus

|  |  | Nucleus |  |  |  |  |  |  |  |
| --- | --- | --- | --- | --- | --- | --- | --- | --- | --- |
|  |  | Nucleus |  | Nucleolus |  | Euchromatin |  | Heterochromatin |  |
| Select of 9 chlorella cells | Number | Volume(μm³) | Area(μm²) | Volume(μm³) | Area(μm²) | Volume(μm³) | Area(μm²) | Volume(μm³) | Area(μm²) |
| Meristem | 1 | 0.88 | 5.28 | 0.012 | 0.27 | 0.063 | 1.91 | 0.28 | 8.96 |
|  | 2 | 0.96 | 5.1 | 0.015 | 0.34 | 0.082 | 2.08 | 0.232 | 8.86 |
|  | 3 | 0.91 | 5.03 | 0.012 | 0.26 | 0.072 | 1.74 | 0.295 | 9.61 |
| Elongation | 4 | 1.16 | 5.96 | 0.032 | 0.61 | 0.098 | 2.97 | 0.27 | 9.62 |
|  | 5 | 1.19 | 6.46 | 0.024 | 0.51 | 0.1 | 3.13 | 0.26 | 9.76 |
|  | 6 | 1.23 | 6.15 | 0.023 | 0.45 | 0.11 | 3.68 | 0.28 | 9.97 |
| Mature | 7 | 1.4 | 6.57 | 0.044 | 0.77 | 0.187 | 6.07 | 0.3 | 11.9 |
|  | 8 | 1.39 | 6.94 | 0.061 | 0.99 | 0.147 | 4.66 | 0.285 | 12.9 |
|  | 9 | 1.34 | 6.83 | 0.032 | 0.52 | 0.132 | 3.71 | 0.282 | 10.9 |

**Table 5:** Volume and area of pyrenoid

|  |  | Pys |  | Py<sub>L</sub> |  | Py (Py<sub>S</sub>+Py<sub>L</sub>) |  |
| --- | --- | --- | --- | --- | --- | --- | --- |
| Select of 9 chlorella cells | Number | Volume(μm³) | Area(μm²) | Volume(μm³) | Area(μm²) | Volume(μm³) | Area(μm²) |
| Meristem | 1 | 0.019 | 0.58 | 0.079 | 1.63 | 0.1 | 2.21 |
|  | 2 | 0.03 | 0.53 | 0.08 | 1.8 | 0.11 | 2.33 |
|  | 3 | 0.02 | 0.46 | 0.08 | 1.8 | 0.1 | 2.26 |
| Elongation | 4 | 0.071 | 1.03 | 0.086 | 1.66 | 0.16 | 2.69 |
|  | 5 | 0.032 | 0.79 | 0.078 | 1.47 | 0.11 | 2.26 |
|  | 6 | 0.097 | 1.18 | 0.138 | 2.8 | 0.23 | 3.98 |
| Mature | 7 | 0.208 | 2.17 | 0.24 | 2.72 | 0.45 | 4.89 |
|  | 8 | 0.21 | 2.24 | 0.23 | 2.6 | 0.44 | 4.84 |
|  | 9 | 0.219 | 2.33 | 0.244 | 2.84 | 0.46 | 5.17 |

**Table 6:** Number and volume of starch grain

| SG volume(μm³) |  |  |  |  |  |  |  |
| --- | --- | --- | --- | --- | --- | --- | --- |
| Number | Meristem | Elongation | Mature | Number | Meristem | Elongation | Mature |
| 1 | 0.030 | 0.030 | 0.033 | 28 |  | 0.140 | 0.103 |
| 2 | 0.030 | 0.043 | 0.035 | 29 |  | 0.140 | 0.105 |
| 3 | 0.031 | 0.045 | 0.031 | 30 |  | 0.140 | 0.110 |
| 4 | 0.047 | 0.048 | 0.038 | 31 |  | 0.141 | 0.111 |
| 5 | 0.069 | 0.048 | 0.038 | 32 |  | 0.161 | 0.120 |
| 6 | 0.081 | 0.048 | 0.048 | 33 |  | 0.168 | 0.121 |
| 7 | 0.085 | 0.053 | 0.048 | 34 |  | 0.178 | 0.123 |
| 8 | 0.085 | 0.055 | 0.048 | 35 |  | 0.218 | 0.131 |
| 9 | 0.086 | 0.056 | 0.050 | 36 |  | 0.225 | 0.142 |
| 10 | 0.096 | 0.066 | 0.050 | 37 |  | 0.291 | 0.143 |
| 11 | 0.099 | 0.072 | 0.051 | 38 |  |  | 0.145 |
| 12 | 0.117 | 0.073 | 0.053 | 39 |  |  | 0.148 |
| 13 | 0.124 | 0.076 | 0.054 | 40 |  |  | 0.153 |
| 14 | 0.144 | 0.096 | 0.055 | 41 |  |  | 0.153 |
| 15 | 0.148 | 0.102 | 0.056 | 42 |  |  | 0.168 |
| 16 | 0.174 | 0.107 | 0.059 | 43 |  |  | 0.169 |
| 17 | 0.175 | 0.108 | 0.061 | 44 |  |  | 0.177 |
| 18 | 0.180 | 0.108 | 0.069 | 45 |  |  | 0.186 |
| 19 | 0.206 | 0.110 | 0.069 | 46 |  |  | 0.194 |
| 20 | 0.245 | 0.110 | 0.070 | 47 |  |  | 0.218 |
| 21 |  | 0.118 | 0.075 | 48 |  |  | 0.230 |
| 22 |  | 0.118 | 0.078 | 49 |  |  | 0.244 |
| 23 |  | 0.120 | 0.080 | 50 |  |  | 0.260 |
| 24 |  | 0.123 | 0.089 | 51 |  |  | 0.271 |
| 25 |  | 0.133 | 0.094 | 52 |  |  | 0.280 |
| 26 |  | 0.136 | 0.095 | 53 |  |  | 0.330 |
| 27 |  | 0.138 | 0.099 | 54 |  |  | 0.380 |

**Table 7:**
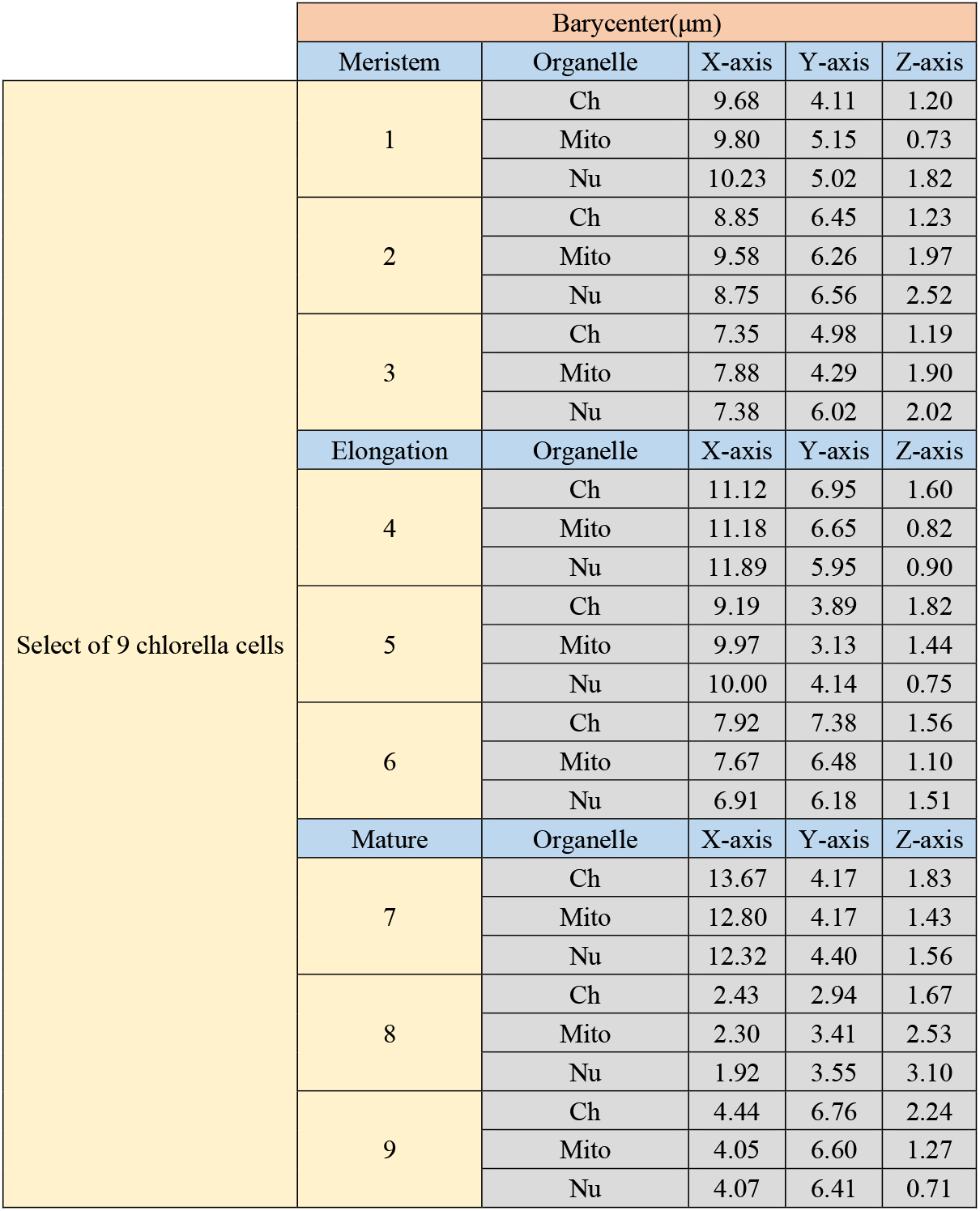
Barycenter of organelles

**Table 8:** D1 and d2 in three growth periods of cell

| d1: the distance of Mito to Ch |  |  |  |  |  |
| --- | --- | --- | --- | --- | --- |
| d2: the distance of Mito to Nu |  |  |  |  |  |
|  |  | d1 | d2 | d1/(d1+d2) | d2/(d1+d2) |
| Meristem | 1 | 1.17 | 1.15 | 0.50 | 0.50 |
|  | 2 | 1.05 | 1.06 | 0.50 | 0.50 |
|  | 3 | 1.81 | 1.13 | 0.62 | 0.38 |
| Elongation | 4 | 1.23 | 1.15 | 0.52 | 0.48 |
|  | 5 | 1.00 | 0.85 | 0.54 | 0.46 |
|  | 6 | 0.91 | 1.05 | 0.46 | 0.54 |
| Mature | 7 | 0.59 | 1.00 | 0.37 | 0.63 |
|  | 8 | 0.55 | 0.95 | 0.37 | 0.63 |
|  | 9 | 0.70 | 0.99 | 0.41 | 0.59 |

**Note (abbreviation for organelle):**

## Acknowledgements

This study was supported by National Key Research and Development Program-China (2017YFE0122800), and Zhejiang Provincial Key Research and Development Program-China (2020C04006). We thank the Center of Cryo-Electron Microscopy (CCEM) at Zhejiang University for technical assistance on FIB-SEM.

## Note (abbreviation for organelle)

CW: cell wall
Ch: chloroplast
Th: thylakoid
SG: starch grain
Py: pyrenoid
Va: vacuole
ER: endoplasmic reticulum
Mito: mitochondrion
Nu: nucleus
LD: lipid droplet
Nuo: nucleolus
Ec: euchromatin
Hc: heterochromatin.

## Notes

### Competing Interest Statement

The authors have declared no competing interest.

